# Mitochondrial Stress Disassembles Nuclear Architecture through Proteolytic Activation of PKCδ and Lamin B1 Phosphorylation in Neuronal Cells: Implications for Pathogenesis of Age-related Neurodegenerative Diseases

**DOI:** 10.1101/2024.11.01.621517

**Authors:** Adhithiya Charli, Jie Luo, Bharathi Palanisamy, Emir Malovic, Zainab Riaz, Cameron Miller, Yuan-Teng Chang, Manikandan Samidurai, Gary Zenitsky, Huajun Jin, Vellareddy Anantharam, Arthi Kanthasamy, Anumantha G Kanthasamy

**Author notes:** Corresponding Author: Anumantha G. Kanthasamy, Department of Physiology and Pharmacology, College of Veterinary Medicine, 500. D.W. Brooks Dr., Coverdell Center, University of Georgia, Athens, GA 30602.

## Abstract

Mitochondrial dysfunction and oxidative stress are hallmarks of pathophysiological processes in age-related neurodegenerative diseases including Parkinson’s, Alzheimer’s and Huntington’s diseases. Neuronal cells are highly vulnerable to mitochondrial stress, however, the cellular and molecular mechanisms underlying the enhanced vulnerability are not well understood. Previously, we demonstrated that the novel PKC isoform PKCδ is highly expressed in dopamin(DA)ergic neurons and plays a key role in inducing apoptotic cell death during neurotoxic stress via caspase-3-mediated proteolytic activation. Herein, we further uncovered a key downstream molecular event of PKCδ signaling following mitochondrial dysfunction that governs neuronal cell death by dissembling nuclear architecture. Exposing N27 DAergic cell line to the mitochondrial complex-1 inhibitor tebufenpyrad induced PKCδ phosphorylation at the T505 activation loop accompanied by caspase-3-dependent proteolytic activation of the kinase. Subcellular analysis using high-resolution 3D confocal microscopy revealed that proteolytically activated cleaved PKCδ translocates to the nuclear compartment, colocalizing with Lamin B1. Electron microscopy also enabled the visualization of nuclear membrane damage triggered by subjecting the DAergic neuronal cells by Tebufenpyrad (Tebu) toxicity. *In silico* analyses identified that the threonine site on Lamin B1 (T575) is likely phosphorylated by PKCδ, suggesting that Lamin B1 serves as a key downstream target of the kinase. Interestingly, N27 DAergic cells stably expressing the PKCδ proteolytic cleavage site-resistant mutant failed to induce nuclear damage, PKCδ activation, and Lamin B1 phosphorylation. Furthermore, CRISPR/Cas9-based stable knockdown of PKCδ greatly attenuated Tebu-induced Lamin B1 phosphorylation. Also, studies using Lamin B1^T575G^ mutated at phosphorylation and PKCδ-ΔNLS-overexpressing N27 cells showed that PKCδ activation and translocation to the nuclear membrane are critically required for phosphorylating Lamin B1 at T575 to induce nuclear membrane damage during Tebu insult. Additionally, Tebu failed to induce Lamin B1 damage and Lamin B1 phosphorylation in organotypic midbrain slices cultured from PKCδ^-/-^ mouse pups. More importantly, we observed higher PKCδ activation, Lamin B1 phosphorylation and Lamin B1 loss in nigral DAergic neurons from the postmortem brains of PD patients as compared to age-matched healthy control brains, thus providing translational relevance of our finding. Collectively, our data reveal that PKCδ functions as a Lamin B1 kinase to disassemble the nuclear membrane during the neuronal cell death process triggered by mitochondrial stress. This mechanistic insight may have important implications for the etiology of age-related neurodegenerative diseases resulting from mitochondrial dysfunction as well as for the development of novel treatment strategies.

## Introduction

Parkinson’s disease (PD) is a chronic progressive neurodegenerative disorder manifested by severe movement difficulties, affecting about 1% of adults older than 60 years. This disease is mainly attributed to the progressive loss of neurons in the substantia nigra (SN), which results in a drastic depletion of striatal dopamine (DA), but its cause is unclear in most individuals (Miller et al., 2021; Poewe et al., 2017; Samii et al., 2004). The main clinical features of PD include akinesia, tremors, rigidity, bradykinesia, walking difficulties, and postural instability as well as non-motor deficits including anosmia, constipation, sleep disorders and other autonomic symptoms (Reinoso et al., 2015; Zis et al., 2015). Among various pathological mechanisms of PD, mitochondrial dysfunction, oxidative stress, neuroinflammation and aberrant protein degradation pathways have been actively pursued. Although the etiology of PD remains enigmatic, growing evidence points to multiple triggers, resulting from interactions between various environmental, genetic, epigenetic and inflammatory factors (Levy et al., 2009; Maiti et al., 2017). Among all these factors, environmental triggers have been closely linked to mitochondrial dysfunction in PD, because several environmental chemicals severally perturb mitochondrial function. In particular, neurotoxic pesticide exposure induces aberrant mitochondrial morphology and changes in their dynamics, leading to enhanced ROS formation, which, in turn, may aggravate or promote a self-perpetuating cycle of mitochondrial dysfunction and oxidative stress (Charli et al., 2016b; Lin et al., 2012; Schapira, 2007).

Chronic exposure to neurotoxic pesticides has also been shown to cause a number of protein modifications and outcomes that are potentially associated with PD (Corrigan et al., 2000; Fleming et al., 1994; Priyadarshi et al., 2000; Spivey, 2011; Tanner et al., 2011). In particular, persistent exposure to mitochondrial complex-1 inhibiting pesticides has been established to result in PD (Greenamyre et al., 2001; Richardson et al., 2019; Sherer et al., 2007; Testa et al., 2005). Indeed, complex-1 inhibiting pesticides such as rotenone and MPTP have become well-studied models of PD and other neurodegenerative diseases (Anantharam et al., 2007; Dauer and Przedborski, 2003; Ghosh et al., 2010; Jackson-Lewis et al., 2012; Kaul et al., 2003; Terron et al., 2018). We recently showed that tebufenpyrad (Tebu), a mitochondrial complex-1 inhibiting acaricide, functions similarly to rotenone in mediating mitochondrial functional and structural damage in DAergic neuronal cells (Charli et al., 2016a). Tebu is generally used in greenhouses to protect ornamental plants, specifically from mites (EPA PC Code-090102 and IRAC - Insecticide Resistance Action Committee).

PKCδ, a member of the novel PKC family, is a well-characterized, redox-sensitive kinase present in various cell types. Oxidative stress-induced activation of the kinase includes membrane translocation of the activated kinase, tyrosine phosphorylation, and proteolysis. Activation occurs in a caspase-3-dependent manner, during which caspase-3 cleaves the native PKCδ (72-74 kDa) into 41-kDa catalytically active and 38-kDa regulatory fragments, thus marking the activation of the kinase (Anantharam et al., 2002; Brodie and Blumberg, 2003; Kanthasamy et al., 2003; Kaul et al., 2003; Ren et al., 2002). Many studies show that proteolytic activation of PKCδ plays a critical role in accentuating the apoptotic cell death process in neurons and various other cell types. PKCδ is also able to potentiate further apoptotic signaling via positive feedback activation of the caspase cascade, thereby the kinase plays a dual function as a mediator and amplifier of apoptosis (Anantharam et al., 2002; Brodie and Blumberg, 2003; Kanthasamy et al., 2003; Kaul et al., 2003; Ren et al., 2002). Interestingly, we found a high expression of PKCδ in nigral DAergic neurons (Zhang et al., 2007). Our previous investigations on the role of PKCδ in DAergic neurodegenerative processes reveal that caspase-3-dependent proteolytic cleavage of PKCδ results in persistent activation of the kinase and mediates the DAergic cell death triggered by a variety of DAergic neurotoxic insults including the Parkinsonian toxicants MPP^+^ and 6-OHDA in both neuronal culture and animal models (Anantharam et al., 2002; Gordon et al., 2012; Hanrott et al., 2006; Hanrott et al., 2008; Harischandra et al., 2015; Harischandra et al., 2014a; Jin et al., 2011; Jin et al., 2014; Kanthasamy et al., 2010; Kanthasamy et al., 2005; Kaul et al., 2003; Latchoumycandane et al., 2011a; Lin et al., 2012). Interestingly, both pharmacological inhibition of PKCδ activity and genetically manipulating proteolytic PKCδ activation confer resistance to neurotoxic agents that mediate apoptosis in DAergic neuronal cells (Latchoumycandane et al., 2011b; Latchoumycandane et al., 2005; Sun et al., 2008), further implicating a critical role for PKCδ activation during oxidative stress-induced DAergic neuronal cell death. We also have previously demonstrated that expression of the catalytic fragment of PKCδ, but not the regulatory fragment, induces apoptosis in DAergic neurons, demonstrating the proteolytic action in DAergic neuronal cell death (Sun et al., 2008). Emerging evidence suggests that differential compartmentalization of activated PKCδ to subcellular organelles may be a critical determinant of its function in signal transduction (Gomel et al., 2007). Nuclear translocation of PKCδ has been shown to regulate the apoptotic pathway by altering the expression of a range of pro- or anti-apoptotic molecules through the phosphorylation of transcription factors (Brodie and Blumberg, 2003; DeVries et al., 2002; Ren et al., 2002). Despite this evidence, the exact interaction between PKCδ and key nuclear target and its role in mitochondrial dysfunction-linked DAergic neurodegenerative processes need to be elucidated.

In this study, we unraveled the pivotal role of PKCδ’s function as a Lamin B1 kinase, wherein it interacts with the key nuclear membrane subunit lamin B1 to initiate nuclear membrane damage in DAergic neurons, thereby leading to apoptotic neuronal cell death. This novel finding mechanistically demonstrates the phosphorylation of Lamin B1 at T575 by PKCδ, which was activated following mitochondrial dysfunction in DAergic neuronal cells. Also, the importance of PKCδ functioning as a Lamin B1 kinase was further confirmed using CRISPR/Cas9-based PKCδ knockdown (KD) and PKCδ cleavage-resistant mutant cell models. Translational evidence was further obtained by using both organotypic brain slice culture and transgenic (Tg) mouse models of PD to demonstrate how PKCδ activation damages Lamin B1 by phosphorylating it at T575. We also confirmed these findings in human postmortem PD brains. Our findings reveal new mechanistic insights into DAergic degeneration as well as identify potential drug targets for prospective small-molecule discovery that could be used in the neuroprotective treatment for PD.

## Results

### Tebu exposure induces increases in caspase-3 activity and proteolytic activation of PKCδ in N27 mesencephalic DAergic neuronal cells

To investigate the role of the PKCδ signaling pathway in mitochondrial complex 1 inhibition results in DAergic neuronal cell death, we treated N27 DAergic neuronal cells with 3 µM of Tebu for 1-3 h and cell lysates were subject to Western blot measurement of PKCδ activation. Tebu induced a time-dependent processing of PKCδ proteolytic cleavage, significant increase in catalytically active and regulatory PKCδ fragments was evident after exposure to Tebu for 3 h (Fig. 1A). Both Western blotting (Fig. 1A) and ICC (Fig. 1B) further reveal that Tebu increased PKCδ-Thr505 activation-loop phosphorylation, which is required for PKCδ kinase activity and serves as a marker of PKCδ activation. Exposure to Tebu for 3 h also significantly induced caspase-3-dependent PKCδ proteolytic activation compared to untreated N27 cells, as evidenced by increase in caspase-3 enzyme activity (Fig. 1C, 1D). Cumulatively, our results suggest that Tebu-induced mitochondrial inhibition triggered caspase-3-dependent proteolytic activation of PKCδ in N27 DAergic neuronal cells.

**Figure 1:**
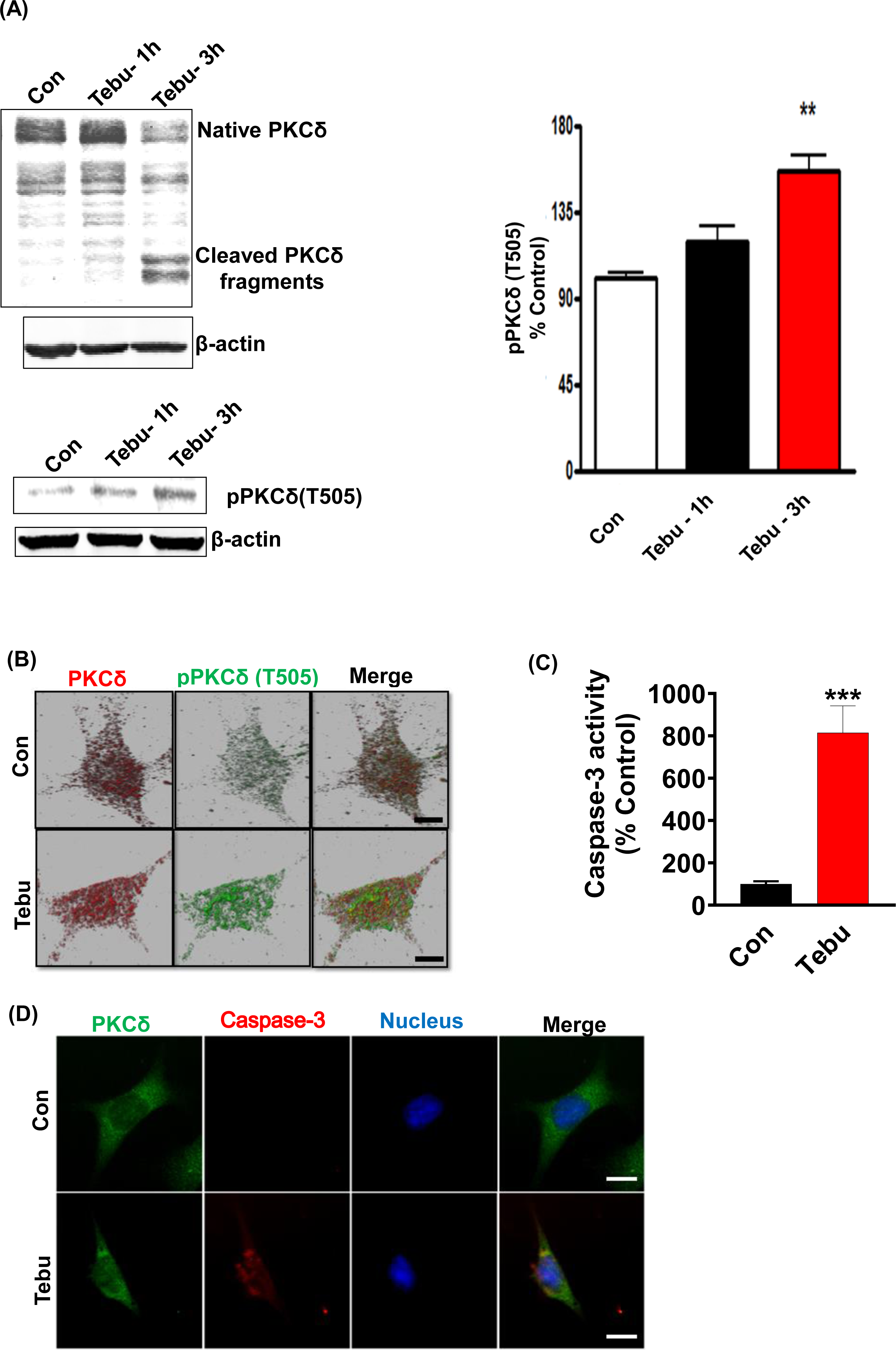
Mitochondrial inhibition triggers PKCδ activation and caspase-3 activity in the DAergic neuronal cell culture model. N27 cells were treated with Tebu (3 µM) for 3 hr. (A) Representative immunoblots of PKCδ and phospho-PKCδ (T505) showing proteolytic cleavage- dependent activation of PKCδ and phosphorylation of PKCδ at T505 induced by Tebu; β-Actin was used as a loading control for all Western blot analyses. (B) Fluorescently stained for PKCδ (red) and phospho-PKCδ at T505 (green). (C) **C**aspase-3 activity assay, using the Ac-DEVD-AMC caspase-3 substrate, showing Tebu-induced caspase-3 activity in N27 cells. (D) Fluorescently stained for Caspase-3 (red) and PKCδ (green) with Hoechst dye staining the nuclei (blue). All the z-stack images in B and D were captured using a 63X oil immersion lens on a Leica confocal microscope. Images were processed using IMARIS 10.0 software. Scale bar, 10 μm. Data shown represent mean ± SEM from two independent experiments performed in triplicate (\*\**p*<0.01 and \*\*\**p*<0.001).

### Proteolytic activated PKC**δ** phosphorylates Lamin B1 to disrupt the nuclear envelope and cause nuclear damage

Next, we investigated the molecular mechanism/pathway underlying PKCδ-mediated DAergic neuron death following exposure to the Tebu. Cross et al, (2000b) reported that PKCδ functioned in the disassembly of the nuclear lamina during the apoptosis. Very recently, a review by Liu et al., (2020) discussed the emerging role in nuclear lamin phosphorylation in gene regulation and pathogenesis of laminopathies (Nucleus. 2020; 11(1): 299–314). This led us to postulate that PKCδ phosphorylates Lamin B1 to induce nuclear damage and subsequent apoptotic DAergic neuronal cell death. To test this possibility, we first performed *in silico* analysis using two online tools to identify potential PKCδ phosphorylation sites in Lamin B1. Using NetPhos 2.0 to predict phosphorylation sites of Lamin B1, we found serine and threonine residues with a probability higher than 95% of being phosphorylated, including the site Threonine 575 (T575) in the XXX motif region (Fig. 2A, left panel). Further analysis using the PhosphoPICK Site analyzer for predicting kinase-specific phosphorylation revealed PKCδ as the most likely kinase to regulate the phosphorylated T575 residue in Lamin B1 (Fig. 2A, right panel). Next, using an available specific antibody raised against the phosphorylated T575 on Lamin B1, we assessed whether Tebu-induced mitochondrial inhibition resulted in phosphorylation of this residue. Western blot analysis showed that exposing N27 cells to 3 µM Tebu for 3 h induced the robust phosphorylation of Lamin B1-T575 residue, which was accompanied by decreased Lamin B1 expression indicating damage to the nuclear membrane (Fig. 2B). Dual immunostaining for Lamin B1 and activated PKCδ (phosphorylated PKCδ-T505) revealed the loss of Lamin B1 architectural integrity of the nuclear envelope as well as co-localized activation of PKCδ in Tebu-treated N27 cells (Fig. 2C). Consistent with this finding, TEM revealed severe degradation of the nuclear membrane in Tebu-treated N27 cells (Fig. 2D). We also performed Duolink PLA, which further demonstrated the physical interaction between PKCδ and Lamin B1 in Tebu-treated N27 cells (Fig. 2E). Collectively, these results suggest that exposure to Tebu induced proteolytically activated PKCδ translocates to nuclear membrane, physically interacts and phosphorylates Lamin B1 at T575 residue to cause nuclear damage. *In silico* prediction for site-specific kinase indicates the T575 residue of Lamin B1 is highly targeted by activated PKCδ.

**Figure 2:**
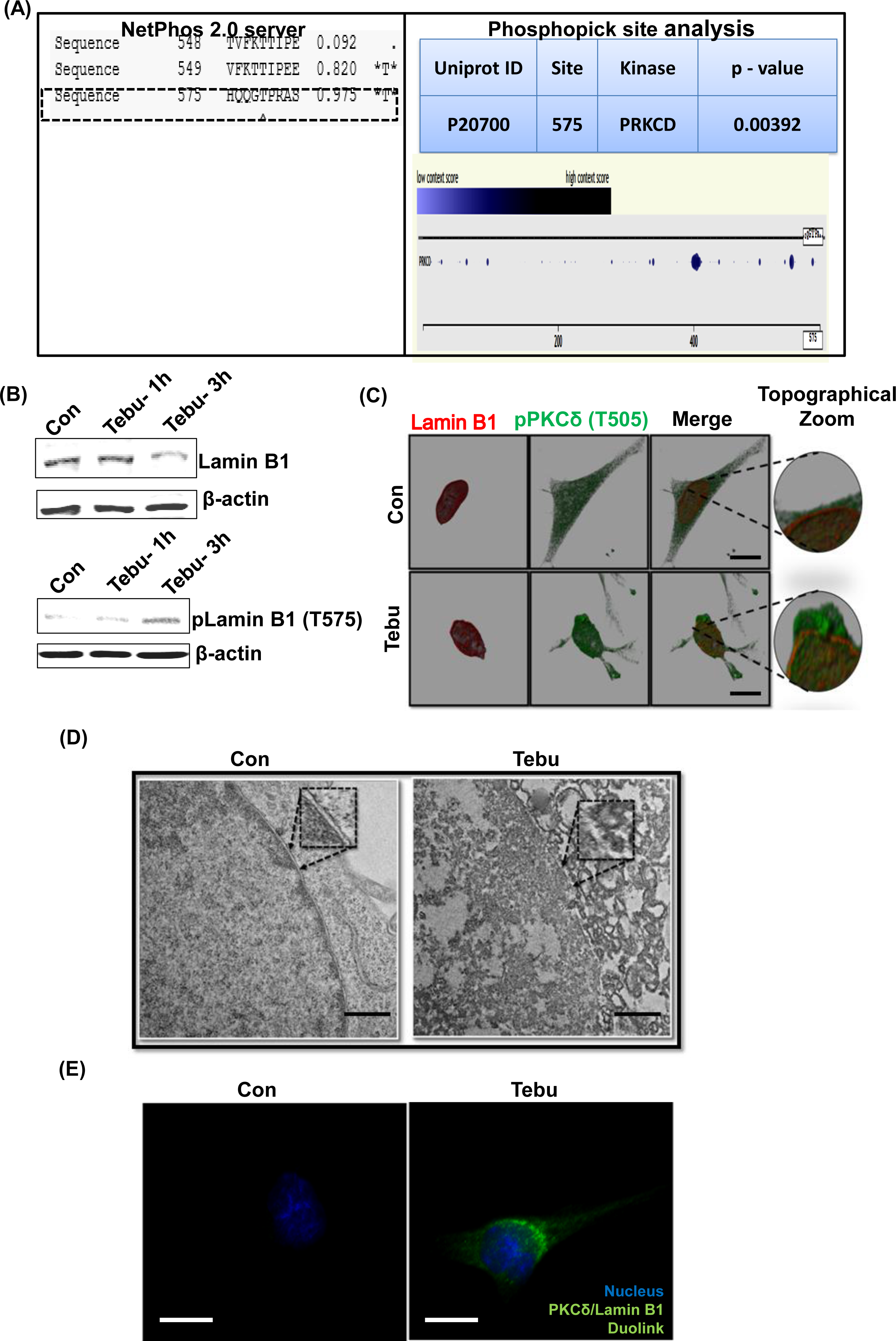
Activated PKCδ functions as a lamin kinase, initiating Lamin B1 phosphorylation, and orchestrates lamin damage. (A) In silico and phospho site-matching analyses using the online tools NetPhos Server 2.0 and PhosphoPICK showing PKCδ site targets on Lamin B1. (B) Representative immunoblots of Lamin B1 and phospho-Lamin B1 (T575) showing that PKCδ phosphorylates Lamin B1 at T575 and induces lamin damage in Tebu-treated N27 cells; β-Actin was used as the Western blot loading control. (C) N27 cells treated with Tebu (3 µM, 3 h) were fluorescently labeled for Lamin B1 (red) and PKCδ (green) and the nuclei were stained Hoechst blue. Scale bar, 10 μm. (D) Representative transmission electron microscopy images depicting the nuclear membrane structure of N27 cells treated with 3 μM Tebu for 3 h. The magnified *inset box*es point to the nuclear envelope structures of control and Tebu-treated cells. Scale bar, 1 μm. (E) Duolink proximal ligation assay reveals that PKCδ and Lamin B1 interact in Tebu-treated N27 cells but not in control cells following 3 h of pesticide exposure. Scale bar, 10 μm.

### Stable CRISPR/Cas9-based PKC**δ** KD or stably expressing PKC**δ** CRM mutant abolishes Tebu-induced Lamin B1-T575 phosphorylation and nuclear membrane damage

To further confirm that T575 is an important phosphorylation site of PKCδ in Lamin B1 during mitochondrial inhibition-associated DAergic neuronal cell death, we examined whether genetically manipulating the PKCδ signaling pathway can interfere with the Tebu-induced Lamin B1-T575 phosphorylation and nuclear membrane damage. For this, we created a stable PKCδ KD N27 cell line by transduction with the CRISPR/Cas9-based PKCδ KD lentivirus. Stable PKCδ KD resulted in a ∼90% loss of PKCδ mRNA levels. CRISPR/Cas9 control and PKCδ KD N27 cells were treated with 3 µM Tebu for 3 h. As expected, CRISPR/Cas9 PKCδ KD cells did not show any detectable PKCδ-Thr505 activation-loop phosphorylation following Tebu exposure (Fig. 3A). Importantly, the PKCδ knockdown almost completely abolished Tebu-induced Lamin B1-T575 phosphorylation as well as the loss of native Lamin B1 (Fig. 3B-C). Confocal ICC analysis of PKCδ and Lamin B1 also revealed resistance to Tebu-induced Lamin B1 loss (nuclear membrane damage) in PKCδ stable KD cells, but not in control cells (Fig. 3D). Next, we tested N27 cell line stably expressing the PKCδ cleavage-resistant mutant (PKCδ-CRM) with 3 µM Tebu for 3 h. No significant PKCδ proteolytic activation or Thr505 activation-loop phosphorylation occurred in Tebu-treated PKCδ-CRM N27 cells (Fig. 3E). Furthermore, overexpressing PKCδ-CRM completely also blocked Tebu-induced Lamin B1 loss and T575 phosphorylation (Fig. 3F). Together, these results further support our hypothesis that proteolytic PKCδ activation plays a critical role in regulating mitochondrial inhibition-induced Lamin B1 T575 phosphorylation in DAergic neuronal cells.

**Figure 3:**
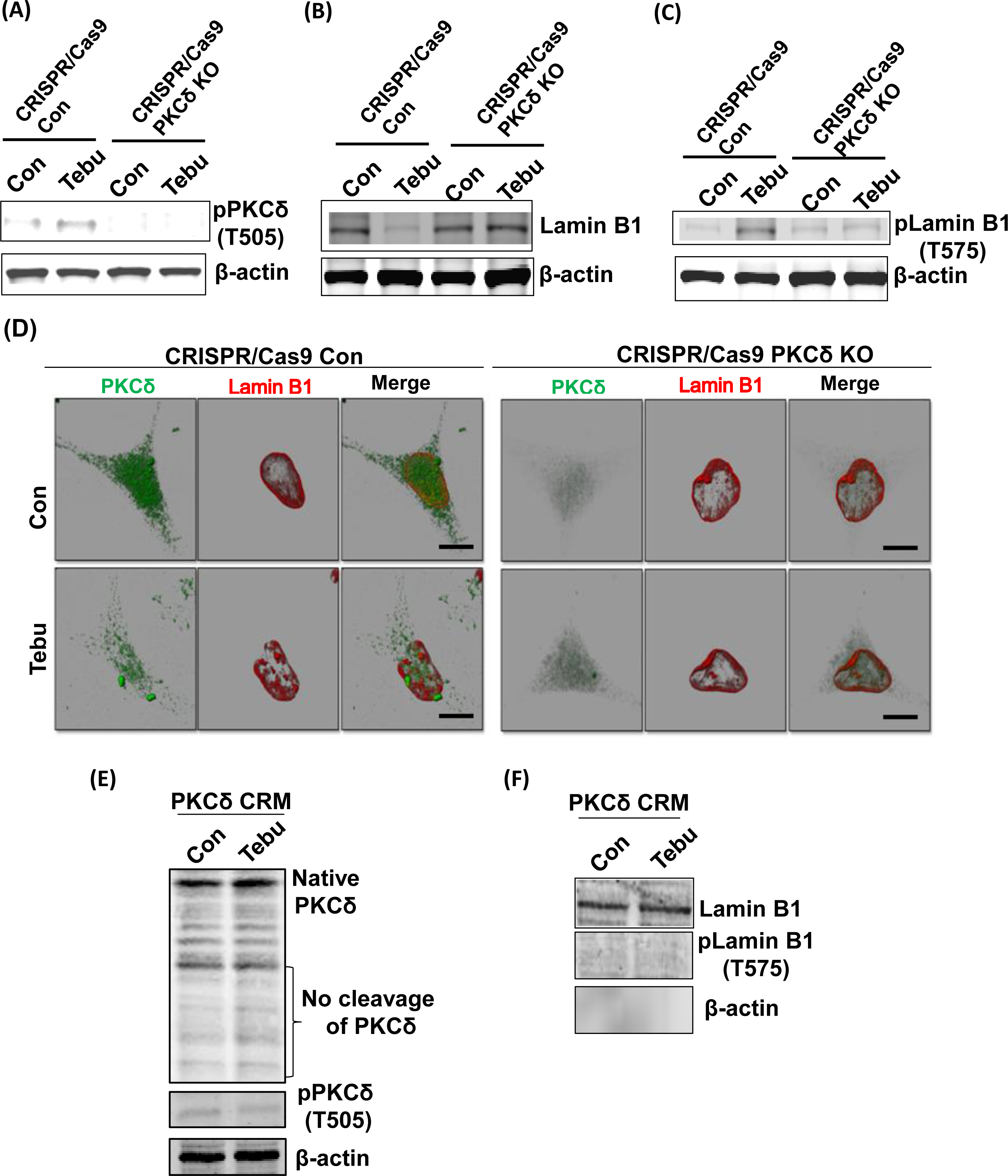
Stable PKCδ CRISPR/Cas9 Knockdown N27 cells and PKCδ cleavage-resistant mutant N27 cells (PKCδ-CRM) were resistant to Tebu-induced PKCδ activation and Lamin B1 phosphorylation. Representative immunoblots of (A) phospho-PKCδ (T505), (B) Lamin B1 and (C) phospho-Lamin B1 (T575) from cells treated with Tebu (3 µM, 3 h); β-Actin was used as loading control for all Western blots. (D) Immunocytochemistry analysis using confocal microscopy showing the resistance of PKCδ CRISPR/Cas9 Knockdown N27 cells to Tebu-induced PKCδ activation and Lamin B1 phosphorylation compared to control CRISPR/Cas9 N27 cells. Scale bar, 10 μm. (E and F) Representative immunoblots depicting the resistance of PKCδ-CRM N27 cells to Tebu-induced (E) PKCδ cleavage and activation and also (F) Lamin B1 phosphorylation and Lamin B1 loss.

### Nuclear translocation of PKC**δ** occurs prior to Lamin B1 phosphorylation

The nuclear translocation of PKCδ is critical for its function in signal transduction. This intracellular translocation process is primarily regulated by the bipartite Nuclear Localization Signal (NLS) present at the C-terminus of PKCδ (between amino acids 611-623). Previous studies from our lab and others have reported that the proteolytic activation of PKCδ leads to the unmasking of the NLS, resulting in the translocation of cleaved constitutively active PKCδ catalytic fragment to the nucleus (DeVries-Seimon et al., 2007; DeVries et al., 2002). To further investigate the mechanism underlying PKCδ-mediated phosphorylation of Lamin B1, two lentiviral plasmid constructs with V5-epitope coding for the NLS deletion PKC^ΔNLS^ mutant was engineered for exogenous expression in N27 cells. Again, treating N27 cells stably expressing PKCδ^WT^with 3 µM of the Tebu for 3 h significantly reduced Lamin B1 and increased phospho-Lamin B1 at T575 compared to untreated cells (Fig. 4A). However, Tebu-treated PKC^ΔNLS^ stably expressing N27 cells demonstrated neither Lamin B1 loss nor Lamin B1 T575 phosphorylation. Furthermore, using confocal microscopy and maximum intensity projection (MIP) imaging, we demonstrated that in PKCδ^WT^N27 stable cells, PKCδ was primarily present in the cytosol and translocated to the nuclear membrane to interact with Lamin B1 upon Tebu exposure. In contrast, stably expressing PKC^ΔNLS^ completely blocked Tebu-induced translocation of PKCδ to the nuclear membrane and subsequent interaction with Lamin B1 (Fig. 4B). This evidence further strengthens our above findings on the role of PKCδ as a Lamin B1 kinase.

**Figure 4:**
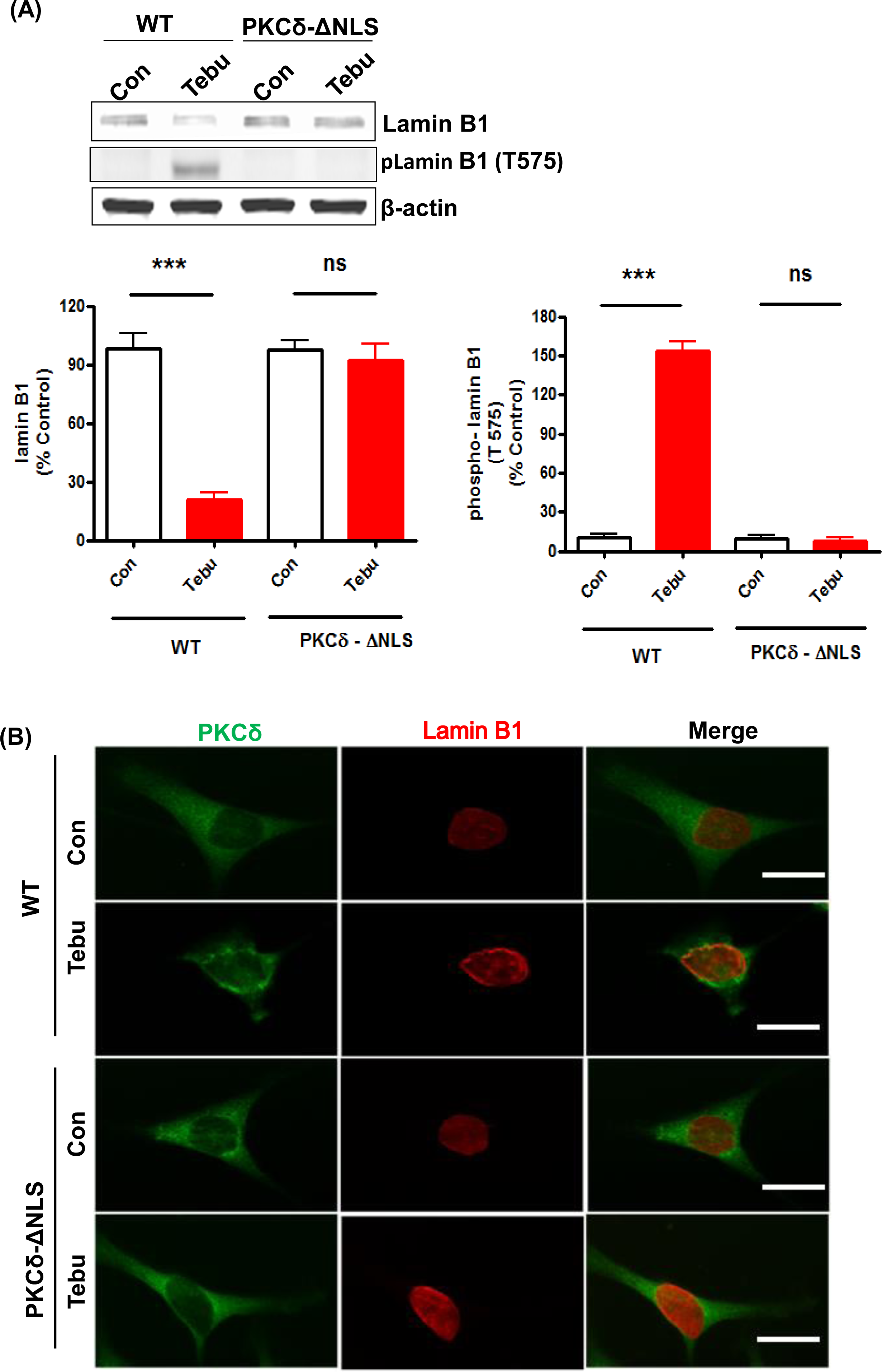
Mutation of the nuclear localization sequence (NLS) of PKCδ prevented its localization with the nuclear membrane post mitochondrial dysfunctional stress and inhibited its functioning as a Lamin B1 kinase. (A) Representative immunoblots (upper panel) and densitometric analyses (lower panels) of Lamin B1 and phospho-Lamin B1 (T575) from wild-type and PKCδ-ΔNLS N27 cells treated with Tebu (3 µM, 3 h); β-Actin was used as the loading control for all Western blots. (B) Immunocytochemistry analysis using confocal microscopy showing the resistance of PKCδ-ΔNLS N27 cells to both Tebu-induced PKCδ translocation to the nuclear membrane and Lamin B1 phosphorylation compared to wild-type N27 cells. Scale bar, 10 μm. Data represent mean ± SEM from two independent experiments performed in triplicate (\*\*\**p*<0.001).

### Expression of a Lamin B1^T575G^ mutant confers resistance to Tebu-induced nuclear membrane damage

To investigate the functional importance of PKCδ-mediated Lamin B1 T575 phosphorylation for -induced nuclear membrane damage, we performed ectopic expression of a Lamin B1^T575G^ mutant in N27 cells. Both native Lamin B1 loss and increased Lamin B1 T575 phosphorylation were observed when Lamin B1^WT^-transfected cells were treated with 3 µM of the Tebu for 3 h (Fig. 5A-B). In contrast, neither Lamin B1 loss or phosphorylation at site T575 occurred in Tebu-treated, Lamin B1^T575^-transfected cells (Fig. 5A-B). This result clearly demonstrates the important role of PKCδ-mediated T575 phosphorylation of Lamin B1, which is required for nuclear membrane damage following Tebu-induced mitochondrial impairment.

**Figure 5:**
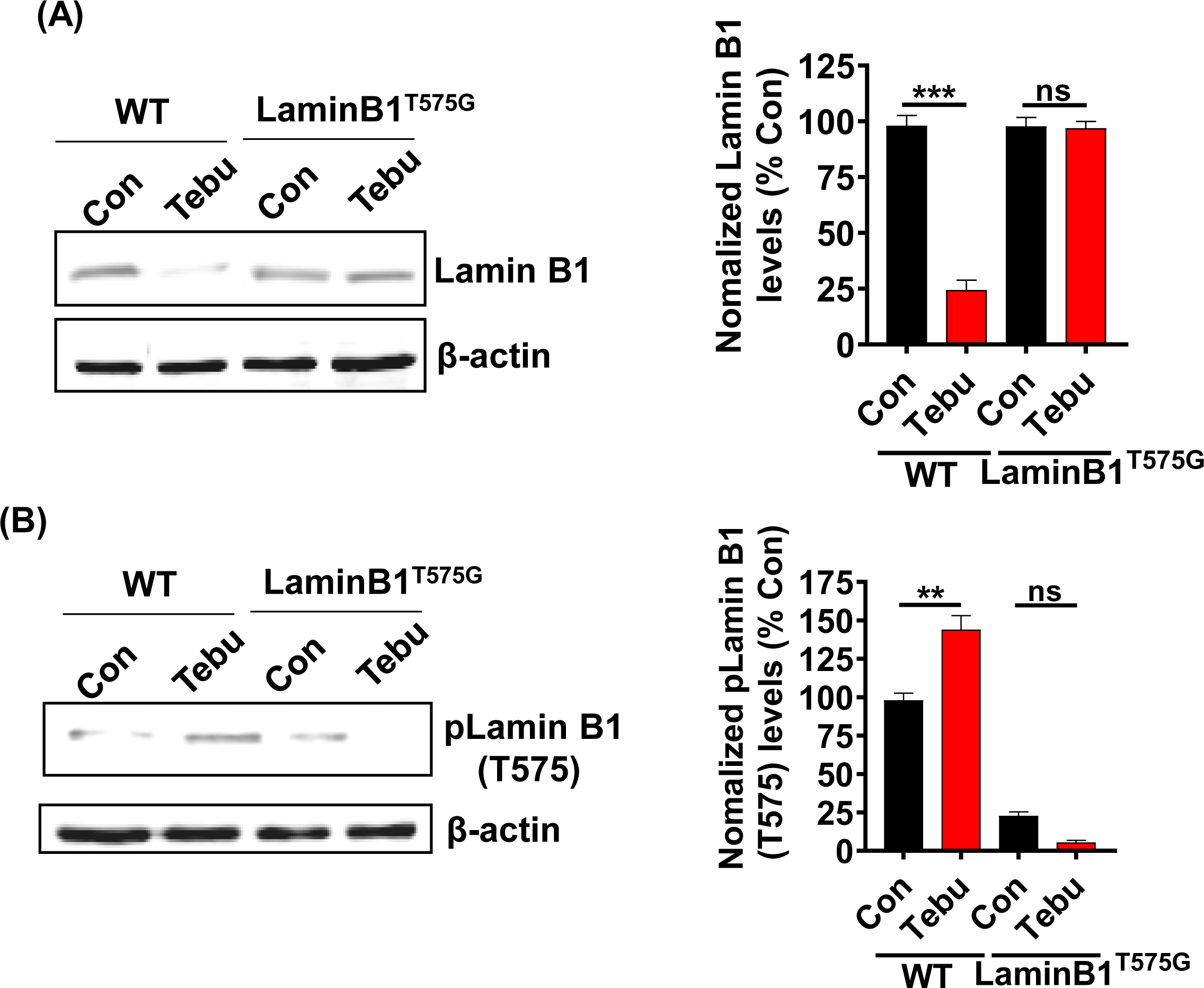
Site-directed mutagenesis of T575 on Lamin B1 prevented its phosphorylation-based activation and damage post-Tebu-induced oxidative stress in N27 dopaminergic neuronal cells. Representative immunoblots and densitometric analyses of (A) Lamin B1 and (B) phospho-Lamin B1 (T575) from wild-type and mutant N27 cells treated with Tebu (3 µM, 3 h); β-Actin was used as a loading control for all Western blots. Data represent mean ± SEM from two independent experiments performed in triplicate (\*\**p*<0.01 and \*\*\**p*<0.001).

### PKC**δ**-deficient organotypic slice cultures demonstrate resistance to Lamin B1 loss and Lamin B1-T575 phosphorylation following exposure to Tebu

Next, we extended our findings obtained from an *in vitro* DAergic cell culture to an *ex vivo* organotypic slice culture model. Organotypic brain slice cultures preserve much of the tissue architecture of the brain region of interest and thereby help validate the translational relevance of mechanistic cell-based toxicity studies (Harischandra et al., 2014b; Humpel, 2015; Kondru et al., 2017; Testa et al., 2005; Ullrich and Humpel, 2009). We treated 350-μm coronal midbrain slices from 9- to 12-day-old PKCδ^+/+^ and PKCδ^-/-^ pups with 20 nM of the Tebu for 24 h. Consistent with our *in vitro* results, mitochondrial inhibition by Tebu induced Lamin B1 loss and T575 phosphorylation in WT slice cultures as evidenced by both Western blot and IHC analyses (Fig. 6A-B). In contrast, PKCδ deficiency significantly diminished Tebu-induced Lamin B1 loss and T575 phosphorylation. These data are consistent with our N27 results, further supporting a functional role for PKCδ in mediating mitochondrial inhibition-associated DAergic cell injury through phosphorylating Lamin B1 and inducing nuclear membrane damage.

**Figure 6:**
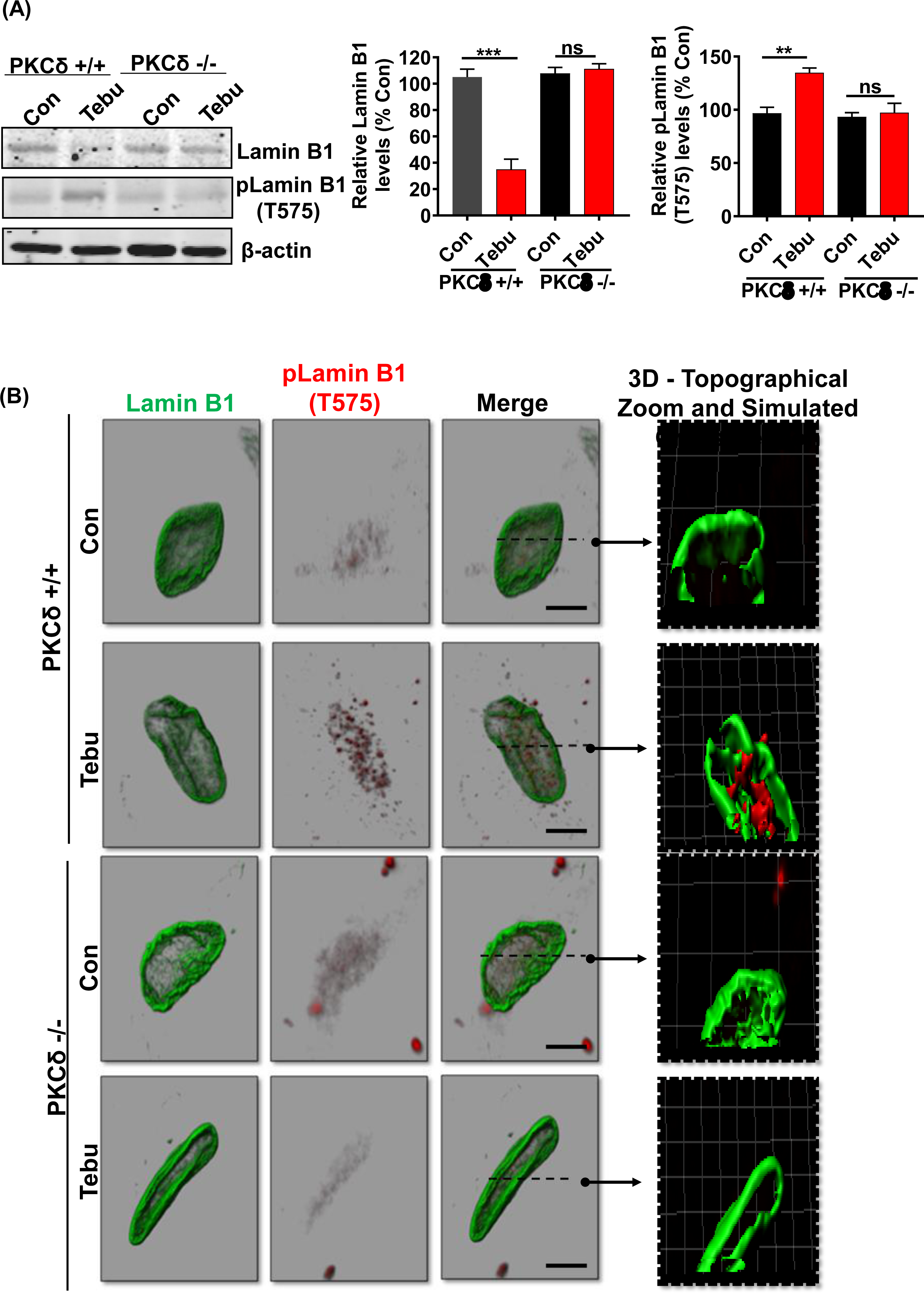
Absence of PKCδ protects against Lamin B1 loss and phosphorylation of Lamin B1 (T575) in organotypic slice cultures. (A) Representative immunoblots of Lamin B1 and p-Lamin B1 (T575) from organotypic PKCδ^+/+^and PKCδ^-/-^ pup brain slices treated with Tebu (20 nM for 24 h); β-Actin was used as the loading control in Western blot analyses. Post treatments the slices were fixed and processed for immunohistochemistry analysis and microscopy. (B) Confocal images depicting fluorescently labeled Lamin B1 (green) and p-Lamin B1 (T575) (red) and Hoechst-stained nuclei (blue). Scale bar, 7.5 μm. Normal shading was retained to more clearly visualize changes in protein expression. Arrows point from simulated cross-section to 3D topographical zoom insets for enhanced visualization of the activated PKCδ in the nuclear envelope.

### PKC**δ** activation and Lamin B1 phosphorylation in a Tg mouse model of mitochondrial dysfunction

We further validated our results in chronic, progressively neurodegenerative Tg MitoPark mice, which were generated by targeted inactivation of the mitochondrial transcription factor A (TFAM) gene in DAergic neurons and exhibit pathological features including mitochondrial dysfunction (Ekstrand et al., 2007) as they relate to human PD. MitoPark mice show significantly decreased locomotion and rearing behavior by age 16 wk when progressive nigrostriatal DAergic degeneration begins (Ekstrand et al., 2007). Western blot revealed significantly reduced levels of Lamin B1 and increased phosphorylation levels of LaminB1-T575 and PKCδ-Thr505 activation-loop phosphorylation in the SN tissues of 20-week-old MitoPark mice as compared to age-matched littermate controls (Fig. 7A). Furthermore, MitoPark SN tissues had significantly lower levels of DA transporter (DAT) relative to littermate controls (Fig. 7A). IHC analysis also depicted enhanced PKCδ-Thr505 activation, Lamin B1-T575 phosphorylation, and loss of Lamin B1 in the SN sections of MitoPark mice (Fig. 7B-C), which were all localized to TH-positive SN DAergic neurons. These results demonstrate PKCδ-mediated Lamin B1-T575 phosphorylation and nuclear membrane damage in a Tg mouse model of mitochondrial dysfunction.

**Figure 7:**
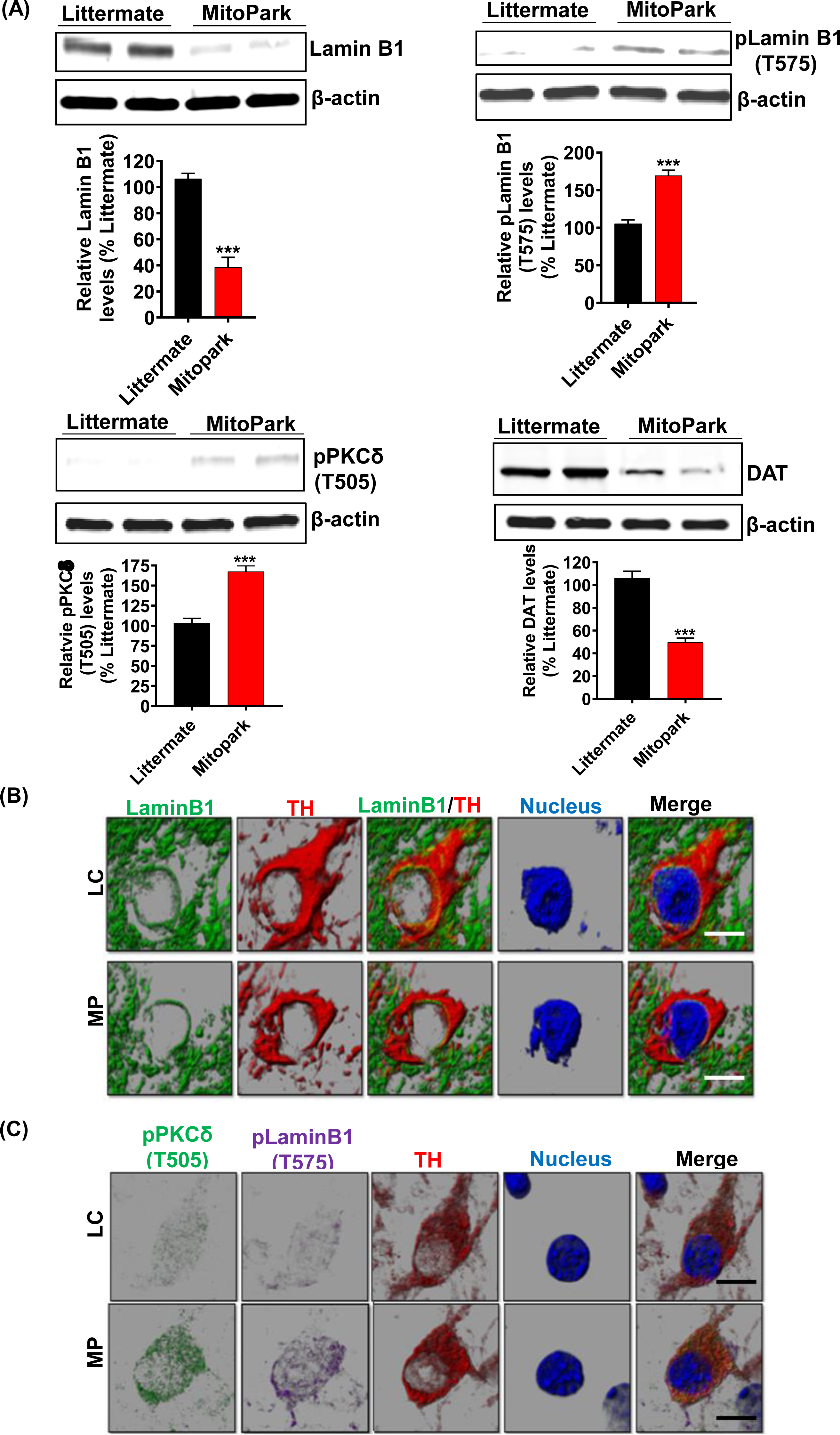
MitoPark transgenic mice demonstrate Lamin B1 loss mediated by activated PKCδ. (A) Representative immunoblots of Lamin B1, p-Lamin B1 (T575), p-PKCδ (T505) and DAT from the *substantia nigra (SN)* of 20-week-old MitoPark and littermate control mice; β-Actin was used as a loading control in Western blots. (B and C) Confocal images of immunohistochemistry performed on 20-week-od MitoPark and littermate SNs showing (B) fluorescently labeled Lamin B1 (green) and tyrosine hydroxylase (TH, red) and Hoechst-stained nuclei (blue) as well as (C) fluorescently labeled p-PKCδ (T505) (green), TH (red) and p-Lamin B1 (T575) (far red) and Hoechst-stained nuclei (blue). Scale bar, 7.5 μm. Normal shading was retained to more clearly visualize changes in protein expression. Data represent mean ± SEM from six mice per group (\*\**p*<0.01).

### Lamin B1 phosphorylation and PKC**δ** activation in the SN of postmortem human PD brains

To further establish the clinical relevance of the PKCδ-Lamin B1 signalling pathway in PD, we evaluated PKCδ activation and Lamin B1 phosphorylation in post-mortem human PD and healthy control SNpc sections and frozen tissues. Western blot analysis revealed a significantly increased levels of phospho-PKCδ (T505) activation in their SN tissues as compared to age-matched control brains (Fig. 8A). Again, this upregulation of PKCδ activation was accompanied by significantly decreased levels of Lamin B1 and higher levels of phospho-Lamin B1 (T575). IHC for Lamin B1 and TH also demonstrated a loss of Lamin B1 expression and nuclear membrane damage in TH-positive DAergic neurons of PD samples as compared to controls (Fig. 8B). Together, these findings demonstrate higher levels of activated PKCδ and Lamin B1 phosphorylation in PD post-mortem tissues compared to age-matched healthy control tissues, further highlighting the important role of both PKCδ and Lamin B1 in the pathogenesis of PD.

**Figure 8:**
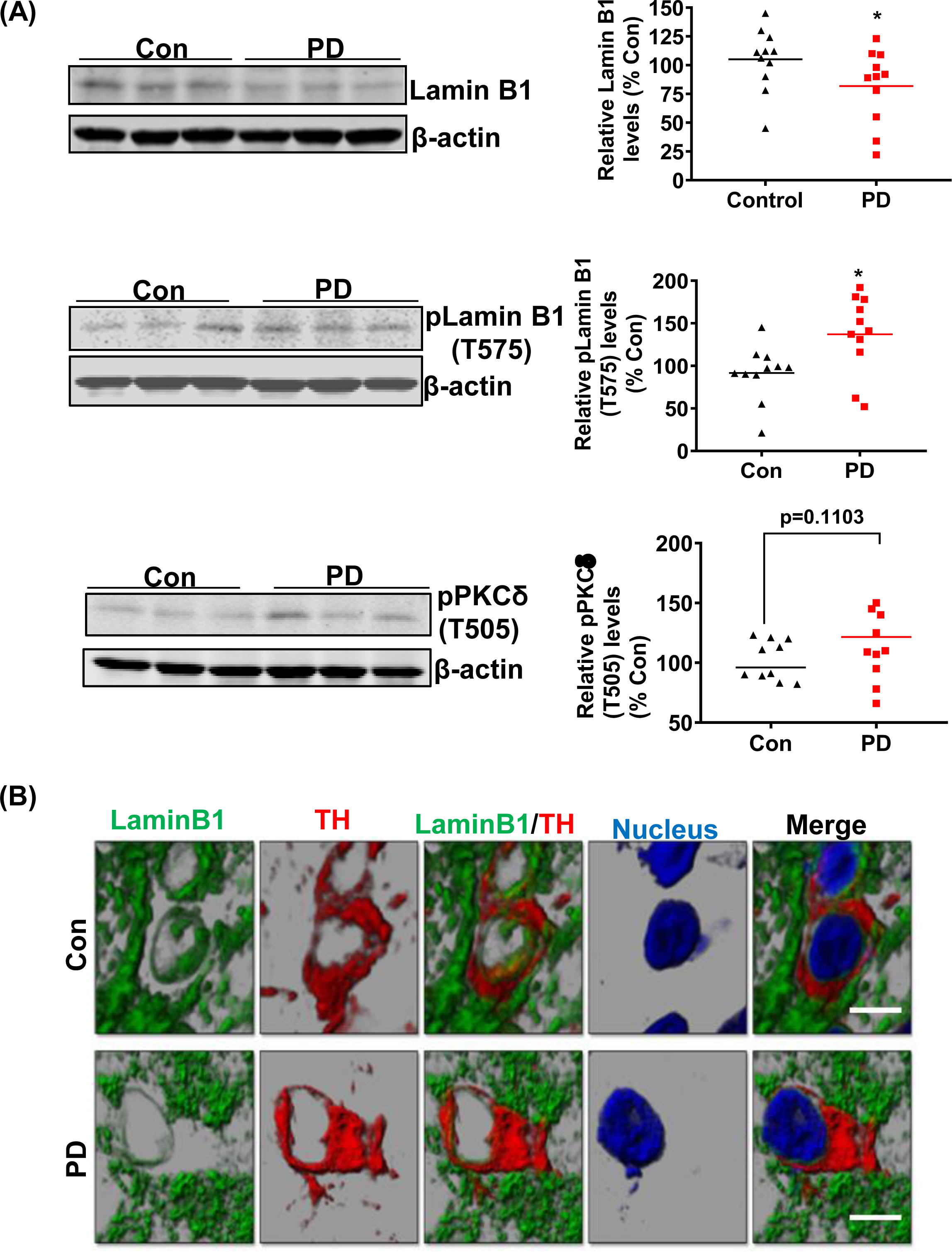
Absence of PKCδ activation, Lamin B1 loss and phosphorylation of Lamin B1 (T575) in the *substantia nigra* (SN) of postmortem PD brains. (A) Representative immunoblots and densitometric analyses of Lamin B1, p-Lamin B1 (T575), and p-PKCδ (T505) from the SN of postmortem control and PD brain tissues obtained from the Miller School of Medicine, University of Miami, FL; β-actin was used as the loading control in Western blot analyses. (B) Confocal images of SN sections from postmortem control and PD brains obtained from Banner Sun Health Research Institute, AZ, showing fluorescently labeled Lamin B1 (green) and TH (red) and Hoechst-stained nuclei (blue). Scale bar, 7.5 μm. Normal shading was retained to more clearly visualize changes in protein expression. Human post-mortem data represent mean ± SEM from 11 control and 11 PD patients (\**p*≤0.05).

## Materials and methods

### Chemicals and reagents

The Tebu (96% purity) was purchased from AK Scientific Inc. (Union City, CA). DMSO and GeneArt Site-directed Mutagenesis System kit (Cat# A13282) were purchased from Fisher Scientific (Fair Lawn, NJ). RPMI 1640 media, fetal bovine serum (FBS), L-glutamine, penicillin, streptomycin pLenti/TOPO expression vector, Fluoro-mount mounting medium (Cat # F4680), Mission Lentiviral Packaging Mix and puromycin were purchased from Sigma-Aldrich (St Louis, MO). The Lenti-X p24 Rapid Titer kit was obtained from Clontech (Cat # 632200, Mountain View, CA). Antibodies for PKCδ, phospho-PKCδ (T505), and Lamin B1 were purchased from Santa Cruz Biotechnology, Inc. (Santa Cruz, CA), while phospho-Lamin B1 (T575) antibody was purchased from Cell Signaling Technology (Danvers, MA). The caspase-3 substrate, Ac-DEVD-AMC, was purchased from Bachem Biosciences, Inc. (King of Prussia, PA).

### Cell culture and treatment paradigm

The rat immortalized mesencephalic DAergic neuronal cell line (1RB3AN27, also known as N27 cells) was a kind gift from Dr. Kedar N. Prasad (University of Colorado Health Sciences Center, Denver, CO). These N27 cells have the potential to differentiate and produce DA in culture when exposed to a suitable cAMP triggering agent, and once the cells are differentiated, they possess increased tyrosine hydroxylase (TH) expression and DA levels (Adams et al., 1996; Zhang et al., 2007). In this study, N27 cells were grown in RPMI 1640 medium containing 10% FBS, 2 mM L-glutamine, 50 units of penicillin, and 50 µg/mL streptomycin, as described previously (Anantharam et al., 2002; Jin et al., 2011c; Prasad et al., 1998). In general, cells plated in a tissue culture plate or flask in accordance with the experimental requirements were cultured overnight in a humidified atmosphere of 5% CO2 at 37°C. The cell density plated for each experiment has been provided in the corresponding method section. The cells were treated with the specified concentration of 3 µM of the Tebu for 0-3 h in serum-free RPMI media. For all experiments with N27 cells, treatments were performed when the cells were 65-70% confluent. Tebu is lipophilic in nature and is hence dissolved in DMSO.

### CRISPR/Cas9-based stable KD of PKCδ in N27 cells

The lentivirus-based CRISPR/Cas9 KD plasmid, pLVU6gRNA-Ef1aPuroCas9GFP-PKCδ, with the PKCδ gRNA target sequence GCGTCGTCCTCCGCCTGCAGGG, was purchased from Sigma-Aldrich. To make lentivirus, the lenti-CRISPR/Cas9 PKCδ KD plasmid and control plasmid were co-transfected into human 293FT cells using the Mission Lentiviral Packaging Mix according to the manufacturer’s instructions. The lentivirus was harvested 48 h post-transfection and titers were measured using the Lenti-X p24 Rapid Titer kit. For stable KD of PKCδ in N27 cells, lentivirus was added into six-well plates containing 0.1 X 106 cells/well at an MOI of 100. After 24 h, fresh media supplemented with puromycin (50 μg/mL) was added to the cells for stable cell selection.

### Establishment of N27 cells stably expressing PKCδ-ΔNLS (nuclear localization signal-deletion mutant), PKCδ-WT, and PKCδ-CRM (cleavage-resistant mutant)

We used the ViraPower Lentiviral gene expression system from Invitrogen (Carlsbad, CA) to generate N27 cell lines stably expressing PKCδ-WT (wild-type; WT), PKCδ-ΔNLS, and PKCδ-CRM. For the preparation of expression vectors containing PKCδ-WT and PKCδ-ΔNLS, 14-2035bp and 14-1843bp of the mPKCδ cDNA were amplified individually with the following primer pairs: CACCATGGCACCCTTCCTGCGC (F) and AATGTCCAGGAATTGCTCAAAC (R) for PKCδ-WT and CACCATGGCACCCTTCCTGCGC (F) and 5’ CTCCAGGAGGGACCAGTT (R) for PKCδ-ΔNLS. All PCR reactions were performed with pfu DNA polymerase to ensure the fidelity of amplification. All PCR products were directly subcloned into the V5-tagged (at the C-terminal) expression vector pLenti/TOPO (Invitrogen) to generate pLenti/PKCδ-WT and pLenti/PKCδ-ΔNLS constructs. To produce lentivirus, pLenti/PKCδ-ΔNLS and pLenti/PKCδ-WT were individually co-transfected with the supporting kit-provided plasmids into 293FT cells by Lipofectamine 2000 as described in the instruction manual. The lentivirus particles in the medium were collected by centrifuging at 3000 rpm for 15 min at 48-72 h post transfection. To generate a stably expressing cell line, lentivirus containing individual pLenti/PKCδ-ΔNLS or pLenti/PKCδ-WT as well as polybrene (6 μg/mL) was added into cultured N27 cells (2 X 105) for 24 h and then replaced with fresh medium. Positive N27 cells were selected by keeping blasticidin (10 μg/mL) in the medium for up to 2 weeks. The lentivirus-based PKCδ-CRM stably expressing N27 cells were created as previously described (Sun et al., 2008).

### Site-directed mutagenesis

To prepare the Lamin B1T575G (threonine to glycine) mutant construct, the mCherry-Lamin B1-10 expression construct (Addgene plasmid # 55069) was used as a template to introduce mutated sequences by using the GeneArt Site-Directed Mutagenesis System kit, according to the manufacturer’s instructions. The synthesis primers used were as follows: GAGGAAGAACTTTTCCACCAGCAGGGAGGCCCAAGAGCATCCAATAGAAGCTG (F) and CACAGCTTCTATTGGATGCTCTTGGGCCTCCCTGCTGGTGGAAAAGTTCTTCCTC (R). The mutant construct was confirmed by DNA sequencing.

### Caspase-3 activity assay

Caspase-3 activity was determined as previously described (Anantharam et al., 2002; Kanthasamy et al., 2008). N27 cells in a density of 1 x 106 cells per T25 cell culture flask were seeded the previous day, and the following day after exposure to the Tebu, the cells were washed once with PBS and re-suspended in lysis buffer containing 50 mM Tris-HCl (pH 7.4), 1 mM EDTA, 10 mM EGTA, and 10 μM digitonin. Cells were then incubated at 37°C for 20–30 min to allow complete lysis. Lysates were quickly centrifuged, and cell-free supernatants were incubated with 50 μM Ac-DEVD-AMC (caspase-3 substrate) at 37°C for 1 h. Caspase activity was then measured using a microplate reader (Molecular Devices Corp., Sunnyvale, CA) with excitation at 380 nm and emission at 460 nm. Caspase activity was calculated as fluorescence units (FU) per mg protein per h and was presented as a percentage of control.

### In silico phosphorylation site analysis

To perform an in silico phosphorylation site analysis on Lamin B1, the web-based program NetPhos 2.0 (http://www.cbs.dtu.dk/services/NetPhos-2.0/) was used to identify the putative serine-threonine phosphorylation sites present in Lamin B1. Further analysis was performed using the online tool PhosphoPICK (http://bioinf.scmb.uq.edu.au/phosphopick/snpanalysis) to identify the kinase(s) that drives site-specific phosphorylation on Lamin B1 as previously described (Patrick et al., 2017; Patrick et al., 2015). Protein sequence data for Lamin B1 and PKCδ were obtained from the Protein Data Bank (PDB).

### Duolink proximal ligation assay (PLA)

Duolink PLA was performed following the manufacturer’s protocol (Thymiakou and Episkopou, 2011) and as previously described (Sarkar et al., 2017). N27 cells were plated on poly-D-lysine (PDL)-coated coverslips at a density of 1 x 104 cells/well in 96-well cell culture plates. Following a 3-h treatment with the Tebu, cells were washed and fixed using 4% paraformaldehyde, blocked with blocking buffer, and incubated in primary antibodies overnight. After primary antibody incubation, the Duolink in situ Detection Reagents Red (Sigma-Aldrich) was added according to the manufacturer’s protocol. Confocal imaging was performed on these coverslips using a DMIRE2 confocal microscope (Leica Microsystems, Buffalo Grove, IL) with a 63X oil objective.

### Organotypic slice culture

Organotypic slice cultures were prepared as previously described (Falsig and Aguzzi, 2008; Harischandra et al., 2014a; Kondru et al., 2017) using whole brains collected from 9-to 12-day-old PKCδ+/+ and PKCδ-/- mouse pups sourced from our Tg PKCδ mouse colony. Brains were oriented inside a specimen tube, prefilled with 2% low-melting-point agarose, for coronal-plane slicing on a vibrating microtome (Compresstome, VF-300, Precisionary Instruments, Greenville, NC). The agarose was quickly solidified by clasping the specimen tube with a chilling block, and then the specimen tube was inserted into the slicing reservoir filled with freshly prepared, ice-cold Gey’s balanced salt solution supplemented with the excitotoxic antagonist, kynurenic acid (GBSSK). The compression lip located in the cutting chamber helps stabilize the brain specimen while obtaining 300-μm thick slices with the blade set at a medium vibration speed. Slices were collected at the specimen tube’s outlet and transferred to another plate prefilled with fresh GBSSK. The slices were then washed twice in 6 mL ice-cold GBSSK, transferred to Millicell 6-well plate inserts (3-5 slices per insert) and were incubated in a humidified atmosphere of 5% CO2 at 37 °C. Culture media was exchanged with fresh media every other day for 10-14 days. Slices were harvested after being treated with the Tebu (20 nM) for 24 h by washing twice in 2 mL of ice-cold PBS.

### MitoPark Animals

Animal procedures were approved and supervised by the Institutional Animal Care and Use Committee (IACUC) at Iowa State University (ISU; Ames, IA). The MitoPark Tg mouse line was originally generated and provided by Dr. Nils-Goran Larson at the Karolinska Institute in Stockholm. The MitoPark mouse model is one in which the mitochondrial transcription factor A (TFAM) is selectively removed in midbrain DA neurons, recapitulates many essential features of PD, such as adult-onset progressive DAergic neurodegeneration, protein aggregation in SN tissues, and L-dopa-responsive motor deficits. TFAM deficiency in this model leads to reduced mitochondrial DNA expression and mitochondrial dysfunction specifically in DAergic neurons, thereby making it a well-established model for understanding the molecular and signaling mechanisms underlying mitochondrial dysfunction in PD (Cong et al., 2016; Ekstrand and Galter, 2009; Ekstrand et al., 2007; Langley et al., 2017). All MitoPark mice used in this study were bred, genotyped, and characterized at ISU. MitoParks and their age-matched littermate control (C57BL/6) animals were maintained under standard conditions of constant temperature (22±1 °C), humidity (relative, 30%), and a 12-h light/dark cycle with free access to food and water as previously described (Ay et al., 2017; Ekstrand et al., 2007; Gordon et al., 2016a; Langley et al., 2017). Equal numbers of male and female mice were randomly assigned to each group. At the age of 20 weeks, mice were sacrificed, and their SN was collected for biochemical and histological studies.

### Human postmortem PD brain samples

We obtained frozen SN tissue samples and cryostat sections from human brains necropsied from patients whose PD was confirmed post-mortem and from age-matched neurologically normal individuals. All human post-mortem samples were procured from brain banks at the Miller School of Medicine, University of Miami, FL, and the Banner Sun Health Research Institute, AZ, and were stored and distributed according to applicable regulations and guidelines involving consent, protection of human subjects, and donor anonymity, and thus Institutional Review Board (IRB) approval from ISU was not required.

### Western blot

Cells or tissues were homogenized and lysed for Western blotting analysis using modified RIPA buffer, as previously described (Kanthasamy et al., 2006; Latchoumycandane et al., 2011). Equal amounts of protein (30-35 μg) were loaded for each sample and separated using 12% SDS-PAGE gels. Proteins were then transferred to a nitrocellulose membrane (Bio-Rad, Hercules, CA) for immunoblotting and blocked using a blocking buffer (Rockland Immunochemicals, Pottstown, PA) specifically formulated for fluorescent Western blotting. The nitrocellulose membrane was incubated overnight with primary antibodies, followed by secondary IR dye-800 conjugated anti-rabbit IgG or Alexa Fluor 680 conjugated anti-mouse IgG for 1 h at room temperature (RT). β-actin was used as a loading control. Western blot images were captured with an Odyssey IR Imaging system (LI-COR. Lincoln, NE) and data were analyzed using Odyssey 3.0 software.

### Immunocytochemistry (ICC)

Immunofluorescence studies in N27 cells were performed according to previously published protocols with some modifications (Gordon et al., 2011; Gordon et al., 2016a). At the end of Tebu treatments, N27 cells grown on PDL-coated coverslips were fixed with 4% paraformaldehyde (PFA), washed in PBS and incubated in blocking buffer (PBS containing 2% BSA, 0.5% Triton X-100 and 0.05% Tween 20) for 1 h at RT. The coverslips were then incubated overnight at 4°C with respective primary antibodies diluted in PBS containing 2% BSA. Samples were then washed several times in PBS and incubated with Alexa Fluor 488 and 555 dye-conjugated secondary antibodies. The nuclei were labeled with Hoechst stain (10 μg/mL) and coverslips were mounted with Fluoromount (F4680) on glass slides for visualization.

### Immunohistochemistry (IHC)

IHC studies were performed on sections from the SN region of MitoPark mice and human post-mortem brain sections as described previously (Ghosh et al., 2013; Gordon et al., 2016a; Jin et al., 2011b; Sarkar et al., 2017). Mice were anesthetized with a mixture of 200 mg/kg ketamine and 20 mg/kg xylazine and then perfused transcardially with freshly prepared 4% PFA and PBS solution. Extracted brains were post-fixed in 4% PFA for 48 h and 30-µm sections were cut using a freezing microtome (CryoStar NX70, Thermo Scientific). Antigen retrieval was performed in citrate buffer (10 mM sodium citrate, pH 8.5) for 30 min at 90°C. Sections were then washed several times in PBS and blocked with PBS containing 2% BSA, 0.2% Triton X-100 and 0.05% Tween 20 for 1 h at RT. Next, sections were incubated with primary antibodies overnight at 4°C and washed 7 times in PBS on a Belly Dancer Shaker (SPI Supplies, West Chester, PA). The sections were then incubated with Alexa Fluor 488, 555, or 663 dye-conjugated secondary antibodies for 75-90 min at RT and their cell nuclei were stained with Hoechst dye. Sections were mounted on slides using Prolong antifade gold mounting medium (Invitrogen) or using the Fluoromount medium according to the manufacturers’ instructions. Samples were finally visualized and analyzed using a Leica DMIRE2 confocal microscope. Similar methods were used for IHC on human postmortem PD brain sections. For mouse slice culture samples, similar IHC methods were employed with minor modifications as previously described (Aguzzi and Falsig, 2012; Harischandra et al., 2014a; Kondru et al., 2017; Sonati et al., 2013).

### Confocal imaging, Z stack image capturing and 3D reconstruction

Confocal imaging was performed at the ISU Microscopy Facility, using a Leica DMIRE2 confocal microscope with a 63X oil objective and Leica Confocal Software. For ICC analysis, one optical series of Z-stack covered 5-7 optical slices representing 0.5-µm thickness each. For IHC analyses of cultured mouse slice sections and human midbrain sections, we employed 12-15 optical slices of 0.5-µm thickness each. IMARIS software 10.0 was used to analyze Z stack images. The surface reconstruction wizard in IMARIS was used to make 3D reconstructed images for seeing architectural changes and enhancing picture definition. Further image details marking topographic alterations were reconstructed using IMARIS modules for normal shading and 3D surface development.

### Transmission Electron Microscopy (TEM)

For TEM, N27 cells grown on coverslips were fixed with 2% glutaraldehyde (w/v) and 2% paraformaldehyde (w/v) in 0.1 M sodium cacodylate buffer, pH 7.2, for 48 h at 4°C. Cells were washed and then fixed in 1% osmium tetroxide in 0.1 M cacodylate buffer for 1 h at RT. The samples were then dehydrated in 70% ethanol, contrast-stained with 2% uranyl acetate in 75% ethanol for 30 min, and further dehydrated in a graded ethanol series. They were then cleared with ultra-pure acetone, infiltrated, and embedded using a modified EPON epoxy resin (Embed 812; Electron Microscopy Sciences, Ft. Washington, PA). Resin blocks were polymerized for 48 h at 70°C. Thick and ultrathin sections were made using a Leica UC6 ultramicrotome. Ultrathin sections were collected onto copper grids and imaged using a JEOL 2100 200kV scanning and transmission electron microscope (Japan Electron Optic Labs USA, Peabody, MA).

### Statistical analysis

All in vitro data were determined from at least 2-3 biologically independent experiments, each done with a minimum of three biological replicates. Prism 4.0 software (GraphPad Software, San Diego, CA) was used to perform one-way ANOVA with the Tukey-Kramer post-test for comparing all treatment groups with that of the control. Differences with p ≤ 0.05 were considered statistically significant.

## Discussion

The current study demonstrates the mechanism of mitochondrial dysfunction by caspase-3-dependent PKCδ proteolytic activation and loss of Lamin. Mitochondrial dysfunction via complex I inhibition has long been implicated as a critical factor contributing to the pathogenesis of diverse acute and chronic neurological disorders, including PD (Schapira et al., 1990), but the molecular mechanisms and the critical regulators of this key pathophysiological hallmark are not well understood. In the present study, we elucidate the role of activated PKCδ in functioning as a Lamin B1 kinase that initiates the process of nuclear membrane damage following mitochondrial dysfunction as demonstrated in N27 DAergic neuronal cultures, animal models of PD, and human postmortem-confirmed PD. Specifically, our study reveals that when mitochondria are inhibited by exposure to the Tebu, the enhanced oxidative stress leads to caspase-3-dependent PKCδ proteolytic activation. Furthermore, the activated PKCδ mediates the phosphorylation of Lamin B1 at T575, thereby leading to Lamin B1 loss and nuclear membrane damage.

Extensive evidence indicates that neurotoxic pesticide exposure can cause mitochondrial dysfunction including abnormalities in oxidative phosphorylation and mitochondrial structure, a decrease in mitochondrial membrane potential, reduced ATP production, and defects in mitochondrial DNA replication, functional dynamics, and respiration. These have all been noted prominently as critical factors contributing to the pathogenesis of PD (Greenamyre et al., 2001; Lin and Beal, 2006; Sherer et al., 2007); (Afeseh Ngwa et al., 2011; Kitazawa et al., 2004; Kitazawa et al., 2001). Our toxicant of interest in this study was Tebu. Classified as a mitochondrial complex 1 inhibitor by the Insecticide Resistance Action Committee (IRAC), Tebu is a greenhouse pesticide used in the USA and many other parts of the world (EPA PC Code-090102) primarily to protect ornamental plants against the action of mites. We previously reported the neurotoxic potential of Tebu in DAergic neuronal cells by demonstrating that exposing N27 cells to Tebu significantly alters mitochondrial functional dynamics and structure (Charli et al., 2016b).

Here, we extend our knowledge by further elucidating the molecular mechanisms underlying Tebu-induced neurotoxicity. In particular, we examined the ability of mitochondrial stress and inhibition following Tebu exposure to initiate PKCδ activation, Lamin B1 loss, and nuclear membrane disassembly. In several of our previous studies, we showed that exposing DAergic neuronal cells to dieldrin and MPP^+^ exacerbates caspase-3-dependent activation of PKCδ, thereby triggering a pro-apoptotic cascade (Kaul et al., 2003; Saminathan et al., 2011; Wiemerslage et al., 2013). Despite this, the major downstream target protein of the PKCδ signaling cascade has yet to be determined. Previously, it was shown that PKCδ functions as an apoptotic lamin kinase in the HL-60 cells (Cross et al., 2000), suggesting the possibility that activated PKCδ may target the phosphorylation of Lamin B1 in DAergic neuronal cells. It is well known that the phosphorylation of lamins disrupts nuclear lamin and orchestrates nuclear membrane damage, especially during mitosis. Moreover, the polymerization of lamins plays a crucial role in maintaining the structural and mechanistic import and export across the nuclear membrane (Machowska et al., 2015; Torvaldson et al., 2015). Our study unravels a novel mechanism by which Lamin B1 is phosphorylated by activated PKCδ during Tebu-stimulated mitochondrial dysfunction in the DAergic neuronal models of PD.

In the present study, the activation of PKCδ in N27 DAergic neuronal cells treated with 3 µM Tebu for 3 h was confirmed by the hallmark proteolytic cleavage and Thr505 activation-loop phosphorylation. Additionally, we observed significantly increased caspase-3 activity in Tebu-treated cells, further supporting a caspase-3-dependent mechanism for PKCδ activation. Following these studies, we systematically identified the specific target of activated PKCδ using the online PhosphoSite analysis tools such as NetPhos Server 2.0 and PhosphoPICK site analyzer. These tools identified Lamin B1 as an interesting PKCδ phosphorylation target. NetPhos predicted the site T575 on Lamin B1 as being the most prone to phosphorylation modification (predicted value - 0.975), while PhosphoPICK further predicted activated PKCδ as the most likely kinase to regulate the phosphorylated T575 residue in Lamin B1. Further analysis using Western blot revealed the downregulation of native Lamin B1 protein and increased phospho-Lamin B1 (T575) levels in Tebu-treated N27 cells. Confocal microscopy confirmed that the activated PKCδ (stained by the Thr505 activation-loop phosphorylation) localized on the nuclear membrane stained with Lamin B1 in the form of a crown-like structure. This co-localization of activated PKCδ and Lamin B1 within the nuclear membrane was further confirmed using Duolink PLA, which also confirmed the physical interaction between PKCδ and Lamin B1 following Tebu-induced mitochondrial stress. Careful inspection of the nuclear membrane using TEM provided us evidence for the loss of Lamin B1, thereby visually confirming nuclear membrane damage in Tebu-exposed N27 cells.

To further strengthen our hypothesis that activated PKCδ serves as a lamin kinase following exposure of N27 cells to Tebu, we generated the PKCδ-CRM stably expressing N27 cells and CRISPR/Cas9-based PKCδ stable KD N27 cells. After Tebu exposure, PKCδ-CRM N27 cells failed to demonstrate either PKCδ activation or any Lamin B1 loss or T575 phosphorylation, providing strong evidence that PKCδ functions as a Lamin B1 kinase mediating Tebu-stimulated nuclear membrane damage. Similarly, the PKCδ CRISPR/Cas9 KD stable N27 cells also did not express any noteworthy Lamin B1 loss or T575 phosphorylation. Immuno-blotting and IHC analysis from the PKCδ-deficient organotypic slice cultures further demonstrated the function of activated PKCδ as a Lamin B1 kinase.

Using the site-directed mutagenesis approach, we highlighted the site-specific phosphorylation of Lamin B1 triggered by PKCδ activation. We created a Lamin B1 mutant construct by replacing the threonine site at 575 with glycine. It was observed that the expression of the Lamin B1^T575G^ mutant did not exhibit any loss of Lamin B1 or phosphorylation of Lamin B1 at site 575 post-Tebu exposure compared to its control. In contrast, the expression of Lamin B1^WT^ did show Lamin B1 loss and T575 phosphorylation. We and others have shown in salivary cells and microglia that the nuclear localization sequence (NLS) located in the C-terminus of PKCδ was responsible for its translocation to the nucleus prior to its activation during oxidative stress conditions (DeVries-Seimon et al., 2007; DeVries et al., 2002; Gordon et al., 2016). In our current study, we employed a mutagenesis technique to mutate the NLS of PKCδ in the N27 cells. Importantly, Tebu treatment was not able to induce Lamin B1 T575 phosphorylation and Lamin B1 loss (nuclear membrane structural impairment) in the PKC^ΔNLS^ of stably expressing N27 cells when compared to the PKCδ^WT^ N27 stable cells.

The MitoPark mouse model of PD is a well-established Tg animal model that has respiratory chain deficiency and mitochondrial dysfunction specifically in DAergic neurons due to the DAergic neuron-specific knockout of TFAM. This model replicates the key hallmark motor and non-motor symptoms as well as the neurochemical changes of PD in humans. Being a well-known Tg animal model for mitochondrial dysfunction, MitoPark mice were utilized to definitively validate the role of activated PKCδ functioning as a Lamin B1 kinase. MitoPark mice at 20 weeks demonstrated significant activation and phosphorylation of PKCδ at T505, substantial loss of native Lamin B1 and phosphorylation of Lamin B1 at T575, compared to the corresponding age-matched littermate controls. The MitoParks also exhibited a drastic reduction in DAT levels compared to age-match controls. Finally, we provide clinical evidence for the functional importance of PKCδ as a Lamin B1 kinase by using human post-mortem SN tissues from PD and control subjects. Overall, our current findings coupled with the other recent observations that substantial loss of LaminB1 in the cellular senescence and induce the mitochondrial dysfunction(Maynard et al., 2022; Saito et al., 2019) The pathological significance of the aforementioned signaling pathways that mitochondrial dysfunction can trigger caspase-3-dependent PKCδ activation is further illustrated by across multiple PD model systems demonstrate, which further translocate to the nuclear membrane and functions as a kinase to phosphorylate Lamin B1 at the T575 residue, resulting in the loss of Lamin B1 and hence damaging the nuclear membrane’s structural integrity followed by the subsequent apoptotic death of the DAergic neurons (Fig. 9A and B). These findings also suggest that Lamin B1 possesses the potential to serve as a promising target for molecular therapy in the treatment of PD.

**Figure 9:**
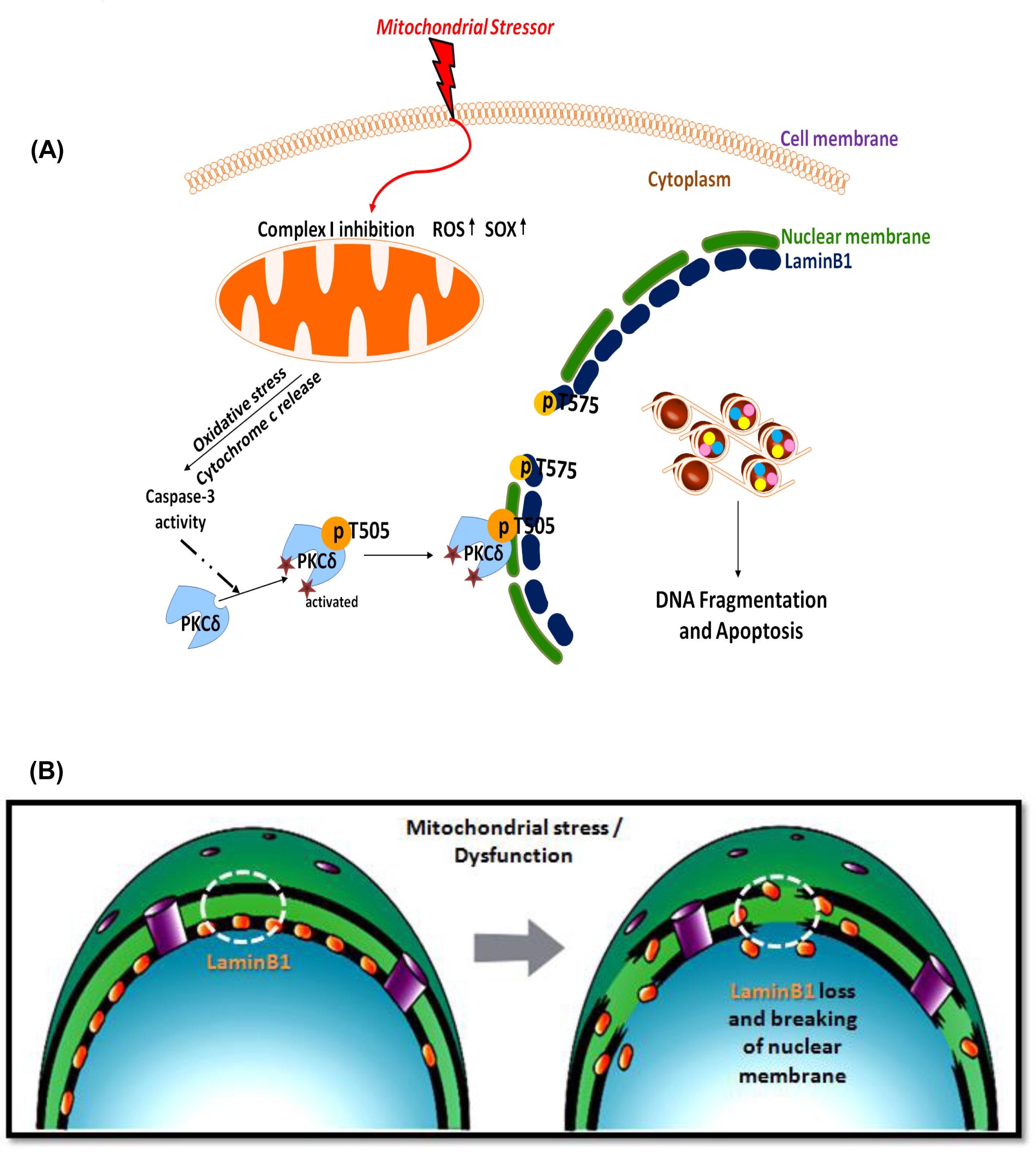
(A) A schematic illustrating the caspase-3-dependent activation of PKCδ mediating phosphorylation of Lamin B1 at T575 and nuclear membrane damage prior to mitochondrial dysfunction in dopaminergic neurons exposed to a mitochondrial stressor. The drawing was created by A. Charli using the Biomedical PowerPoint Toolkit from Motifolio. (B) A schematic illustrating Lamin B1 loss and nuclear membrane damage that occurs following mitochondrial stress.

## Acknowledgments

This work was supported by the National Institutes of Health (NIH) R01 grants ES010586, R01 ES027245, and R01 ES034196. Other sources include W. Eugene and Linda Lloyd Endowed Chair, Johnny Isakson Endowment and the Coach Mark Richt Neurological Disease Research Fund to AGK.

## Potential Conflicts of Interest

All authors declare no actual or potential competing financial interests. A.G.K. has an equity interest in Probiome Therapeutics. The terms of this arrangement have been reviewed and approved by the University of Georgia in accordance with their conflict-of-interest policies.

## Notes

### Competing Interest Statement

The authors have declared no competing interest.

